# Prediction of Adeno-Associated Virus Fitness with a Protein Language Based Machine Learning Model

**DOI:** 10.1101/2024.08.19.608620

**Authors:** Jason Wu, Yu Qiu, Eugenia Lyashenko, Christian Mueller, Sourav R. Choudhury

**Author notes:** These authors contributed equally to this work (Jason Wu, Yu Qiu).

## Abstract

Adeno-associated viral (AAV)-based therapeutics have the potential to transform the lives of patients by delivering one-time treatments for a variety of diseases. However, a critical challenge to their widespread adoption and distribution is the high cost of goods (CoGS). Reducing manufacturing costs by developing AAV capsids with improved yield, or fitness, is key to making gene therapies more affordable. AAV fitness is largely determined by the amino acid sequence of the capsid, however, engineered AAVs are rarely optimized for manufacturability. Here, we report a state-of-the art machine learning (ML) model that predicts the fitness of AAV capsid mutants based on the amino acid sequence of the capsid monomer. By combining a protein language model (PLM) and classical ML techniques, our model achieved a significantly high prediction accuracy (Pearson correlation = 0.818) for capsid fitness. Importantly, tests on completely independent datasets showed robustness and generalizability of our model, even for multi-mutant AAV capsids. Our accurate ML-based model can be used as a surrogate for laborious in vitro experiments, thus saving time and resources, and can be deployed to increase fitness of clinical AAV capsids to make gene therapies economically viable for patients.

## Introduction

The adeno-associated viral (AAV) capsid is composed of three structural proteins: VP1 (87 kDa), VP2 (72 kDa), and VP3 (63 kDa),(1) all encoded by overlapping transcripts of the *cap* gene. Tropism, biodistribution, and capsid assembly are all determined by the capsid sequence. Capsid engineering efforts aim to modify the sequence to improve specificity, biodistribution and safety. However, many capsid sequence changes are deleterious, reducing the fitness of the capsid, and resulting in non-assembling or low-yield capsids.

Despite its potential, the widespread adoption of AAV-based gene therapies faces notable challenges, among them, the high cost of goods. While early AAV gene therapies(2) were based on naturally occurring AAV serotypes, current work aims to enhance efficacy, and safety, and to reduce costs by developing engineered capsids with improved tissue tropism and specificity.(3,4) AAV engineering studies frequently proceed by screening capsid libraries generated by methods such as peptide insertion or saturation mutagenesis of capsid regions. While this approach has yielded many improved capsids, an exhaustive search across the entire capsid sequence could uncover beneficial properties in previously uncharacterized regions of AAV. However, due to the vast search space—20^735^ possible substitutions for AAV2 VP1—such a search not feasible using traditional library screening techniques. Computational methods, including machine learning (ML) approaches, have been successfully applied in drug discovery. For example, virtual screening (VS) predicts interactions between small molecules and targets across the vast landscape of chemicals and proteins.(5) We reasoned that a similar computational approach could predict the fitness of novel capsid sequences, thereby unlocking new capsid regions for exploration.

Machine learning comprises an ensemble of algorithms that can be trained on data sets to create models capable of generalizing to unseen data. One of the important early applications of ML was in natural language processing (NLP), where statistical models were developed for computers to understand and generate human languages. Protein language models (PLMs) were adapted from NLP and have had success in protein structure prediction and protein engineering. (6-8) PLMs treat protein sequences as sentences in which amino acid residues represent words. These models are trained on large protein sequence datasets, learning the “grammar” and “semantics” of protein. The resulting PLMs represent protein sequences as embeddings or numerical representations (embeddings) that capture functional and structural properties of the protein. Two such models are PLM(6) and ESM-2.(9) These models, both trained on the UniRef50 dataset,(10) have markedly different architectures. ESM uses a transformer architecture based on a multi-head attention mechanism. It has no recurrent units and is trained to predict the masked amino acids given the surrounding amino acids. In contrast, PLM uses a multiplicative short-term memory network (mLSTM) based on a recurrent neural network architecture for sequence modeling, combining the long short-term memory (LSTM) and multiplicative recurrent neural network (mRNN) architectures. Unlike ESM-2, PLM is trained to predict the next amino acid in each sequence. Both models have been shown to encode structural and biologically meaningful information about proteins but, because of their different architectures, they may capture different protein properties.(11) Previous studies have used classical ML to predict capsid fitness (a score which captures multiple facets of AAV quality(12)) or assembly, both of which can be used to reduce the number of non-viable AAV variants in libraries(13). We reasoned that the addition of a PLM could complement other ML approaches, taking advantage of additional information about protein structure and function.

Here, we present a method for predicting the fitness of AAV2 mutants based on the amino acid sequence of the capsid monomer VP1 which contains all of the sequence information for the VP1, VP2, and VP3 gene products. Our custom model architecture is trained on a publicly sourced dataset(14) and leverages an ensemble of models using both protein language model and one-hot encoded embeddings. The result is a robust prediction model for AAV fitness. We validated our model on independent datasets and demonstrated that its predictions translate effectively to capsids with mutations across the AAV2 capsid sequence.

## Results

### Model development and training

CAP-PLM is a discriminative model trained to predict the fitness of an AAV2 single-point mutant from the amino acid sequence alone (Figure 1A). Building the CAP-PLM model is broken down into five steps (Supplementary Table 1): reformatting the data, rebalancing the input data embedding the sequences, training a regression model, and predicting fitness of a held-out validation set. For steps 2-4, we evaluated the effect of varying various hyperparameters on the model’s performance. To train and assess the performance of different capsid fitness models, we used publicly available AAV2 fitness data.(14) This library is the most comprehensive AAV capsid mutagenesis library available, comprising fitness scores from a pooled NGS library for all single-point mutations spanning the entire length of the AAV2 *cap* sequence and including all possible single-point insertions, deletions, substitutions, and stop codon mutations. Because this dataset spans the entire *cap* gene rather than focusing on a narrow range,(3) it allows us to predict the effect of mutations across the entire gene. We used the same definition given in the dataset for the fitness score (Supplementary Equation 1). We define the fitness score of a particular capsid as the log2 enrichment of the virus against the input plasmid normalized to parental AAV2 log2 enrichment. With this definition, the fitness score of the wild-type AAV2 is 0, and a fitness score of 1 means that a specific mutant produces twice more virus than AAV2 given the same amount of input plasmid.

**Figure 1.**
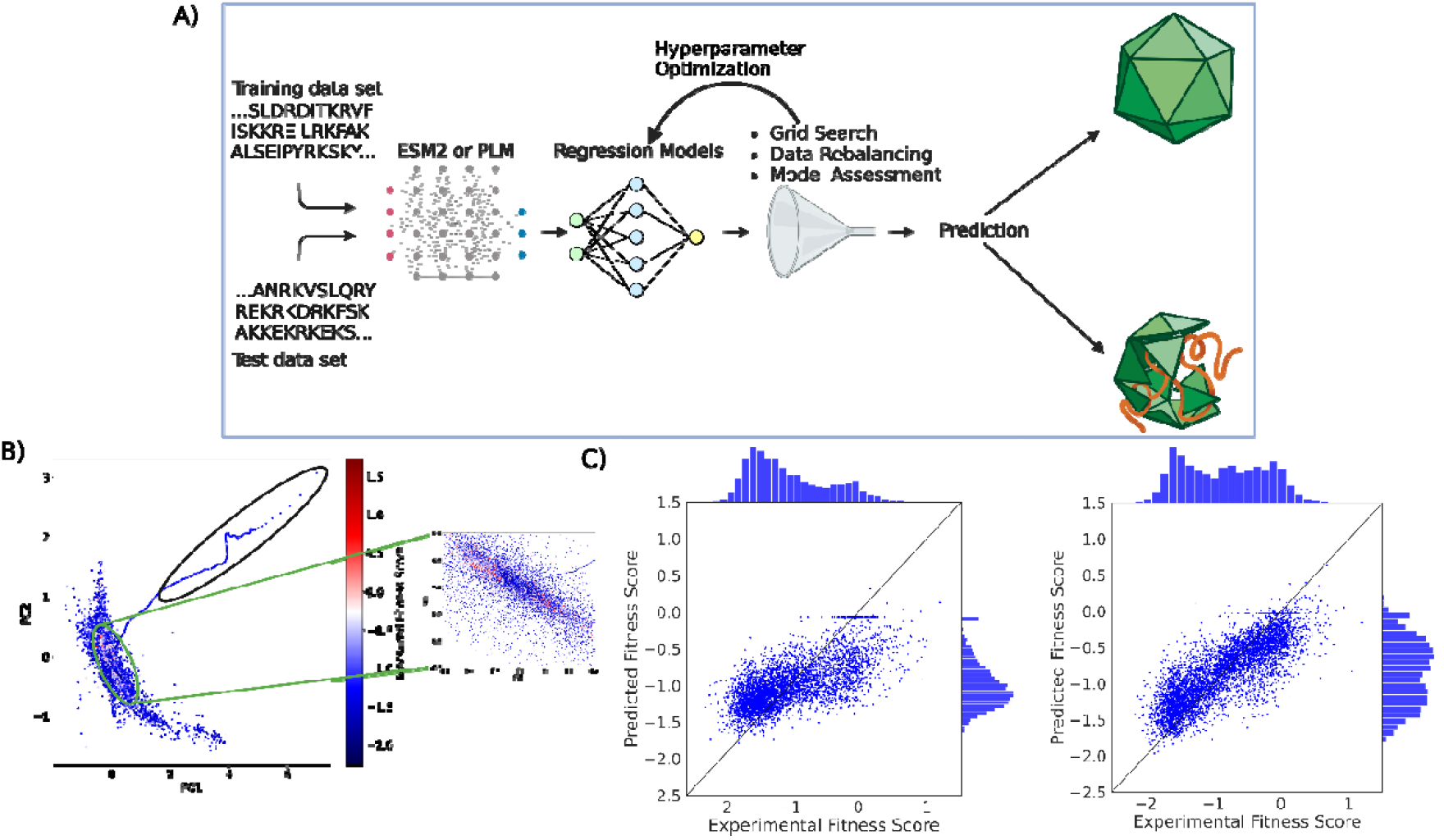
The CAP-PLM model predicts fitness of AAV2 mutants. **(A) The** methodology for training the CAP-PLM fitness prediction model. A basic framework of the different steps taken to turn the AAV2 mutant sequences into usable data for training and prediction is shown above. Each protein sequence is turned into a PLM embedding which is used as the training data for a regression model. This regression model is then validated upon the set aside validation data set to measure model performance. (**B)** PCA on PLM embeddings of VP3. Principal component analysis on PLM embeddings of VPS clusters both low fitness score stop-codon mutations near the beginning of the sequence (black circle) and high fitness score sequences (red circle). (**C)** Comparison between initial and final best model. Pearson correlation of the predictions between the initial and final model increased from 0.678 to 0.818.

Protein embeddings are vector representations encoding structural and functional properties of encodings. PLM’s embeddings have been shown to encode a variety of properties, including structural, physiochemical, and evolutionary information.(6) To show that PLM captures meaningful biological properties of the AAV2 sequence, we conducted Principal Component Analysis (PCA) on the embeddings of AAV2 variants, generated by PLM. We found unsupervised clustering trends related to the fitness of the VP3 region of the AAV2 mutants (Figure 1B). Specifically, we found that the PCA representations of the PLM embeddings cluster high-fitness score mutants together and display a distinctive arm containing stop-codon mutants near the beginning of VP3. Since PLM effectively captures unsupervised biological relationships in the AAV2 single-point mutation dataset, we chose to use PLM embeddings for our downstream model development.

To train our model, we randomized the input dataset, setting aside 20% as the validation data and used the remaining 80% as training data. After training, we tested the model’s ability to predict the fitness of the held-aside test set by calculating Pearson correlation between the prediction and the experimental data (Figure 1C), iteratively testing the effect of varying hyperparameters on model performance. Based on the results of our PCA (Figure 1B), we began by exploring removing stop codon sequences from the input data for model training, but this did not improve model performance (Supplementary Figure 1). To determine whether the VP3 sequence, which comprises a subsection of the VP1 sequence and roughly 5/6 of the proteins in a full capsid,(15) encodes comparable information to the VP1 sequence, we compared the model performance trained by using the full VP1 sequences to that of a model trained using only the VP3 sequences. The result showed 6% improvement on Pearson Correlation of the predicted fitness against the experimental fitness by using VP1 sequences (Supplementary Figure 1; Pearson 0.721 vs 0.678). As 55% of the fitness scores of the input data fell between -0.5 and - 1.5, we reasoned that the model would be likely to overfit on low fitness scores. To combat this, we rebalanced the input data by randomly removing 5095 sequences with scores between -0.5 and -1.5 (about 1/6^th^ of the total). This further improved the model by 5% (Supplementary Figure 1; Pearson correlation of .755 vs .721). Then we tested different regression models, including Ridge, K-Nearest Neighbors, Radius Nearest Neighbors, Support Vector Regression, Nu Support Vector Regression, Random Forest Regression, and Decision Tree Regression, and found that a Random Forest Regression model outperforms others (Supplementary Figure 2).

Next, we explored whether further innovative customization of model architecture can improve the model’s predictive power. A study has reported improving protein fitness prediction by combining a regression model trained on site-specific amino acid features with evolutionary data(16). One-hot encoding is another common way to encode protein sequences. With one-hot encoding, amino acids are each represented as a binary vector, thereby representing the categorial variable of an amino acid as a numerical value. PLMs embeddings can encode information about functional and structural properties of a protein, but one-hot encoding provides only positional properties. Nevertheless, we hypothesized that combining the two might improve the representation. To determine if a composite model trained on both PLM and one-hot encoded embeddings could outperform one trained on PLM embeddings alone, we used the best-performing Random Forest model to predict the fitness scores for the training set and used the differences between this prediction and the ground truth as labels for a second training scheme using one-hot encoded sequences as input. This improved the Pearson correlation by another 8% (Supplementary Figure 1; Pearson correlation of 0.755 to 0.818). Overall, we saw a 20% increase in Pearson correlation over the course of these hyperparameter improvements.

Finally, we evaluated the performance of two pre-trained PLMs PLM and ESM-2 with a downstream model on the fitness prediction task. Interestingly, although transformers in principle capture global dependencies better through self-attention mechanisms, we found that the PLM trained model had a slight advantage in Pearson correlation of 2% over the ESM-2 trained model in this case, in line with observations from other use cases(17) (Supplementary Figure 1; Pearson correlation of .818 vs. .805). Therefore, our final model (Figure 2D) comprises two Random Forest models trained on PLM and one-hot encoded embeddings respectively. To allow for large insertions into the VP1 sequences, we permitted an input of up to 755 amino acids to CAP-PLM.

**Figure 2.**
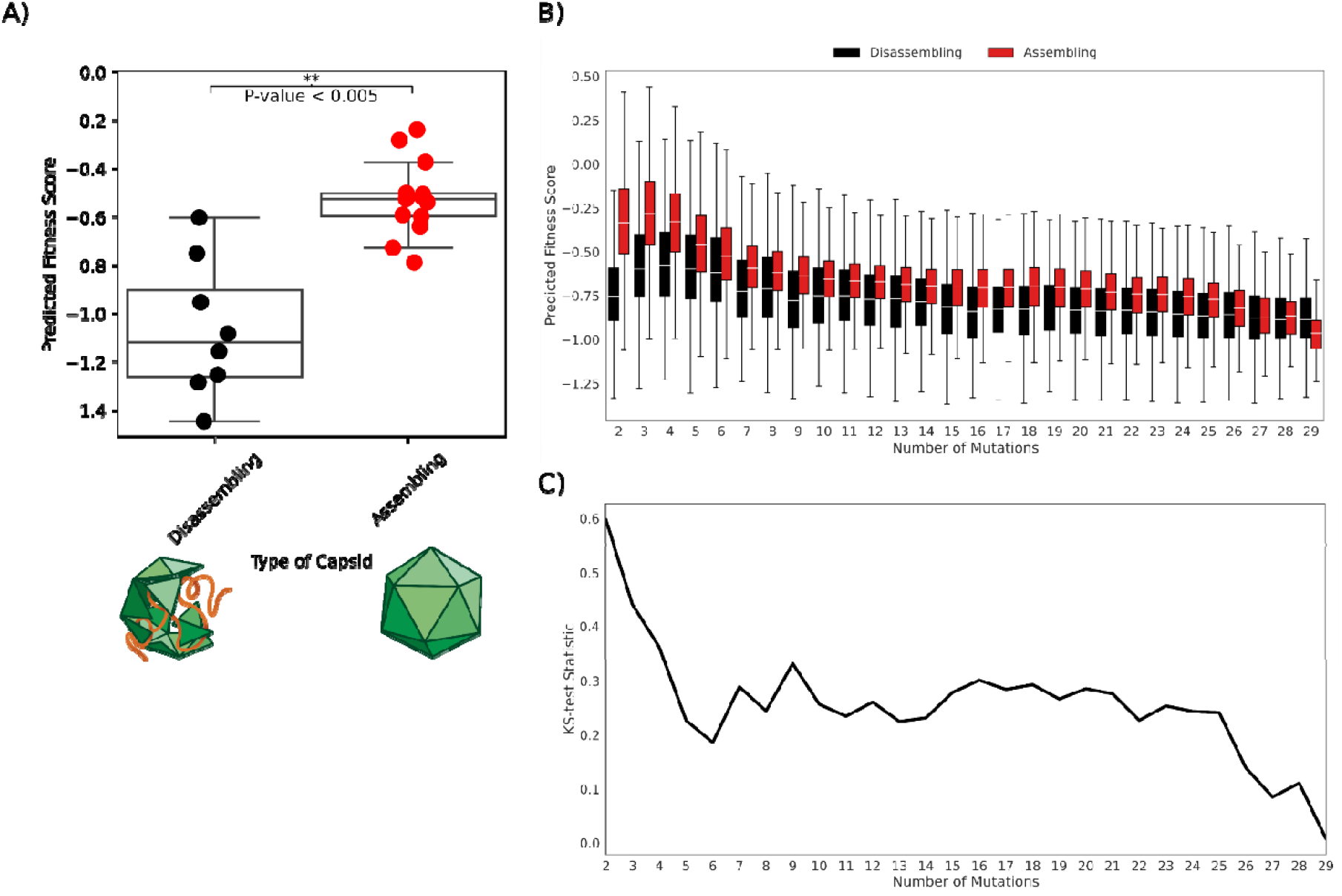
CAP-PLM generalizes to predictions on fitness of AAV2 multiple-mutation mutants. (**A**) By using CAP-PLM to predict the fitness score of 12 different AAV2 mutants, we can distinguish between assembling and non-assembling capsids with more than one mutation away from WT AAV2 using KS-Test. (**B**) Comparison between predicted fitness scores of a 561-588 substitution library. A boxplot of predictions upon multi-mutants of AAV2. We use KS-test to compare the difference. (**C**) Comparison between KS-test statistic of a 561-588 substitution library. A line plot of the KS-test statistic when comparing the predicted fitnes scores of viable vs non-viable mutants with the same number of mutations.

**Figure 3.**
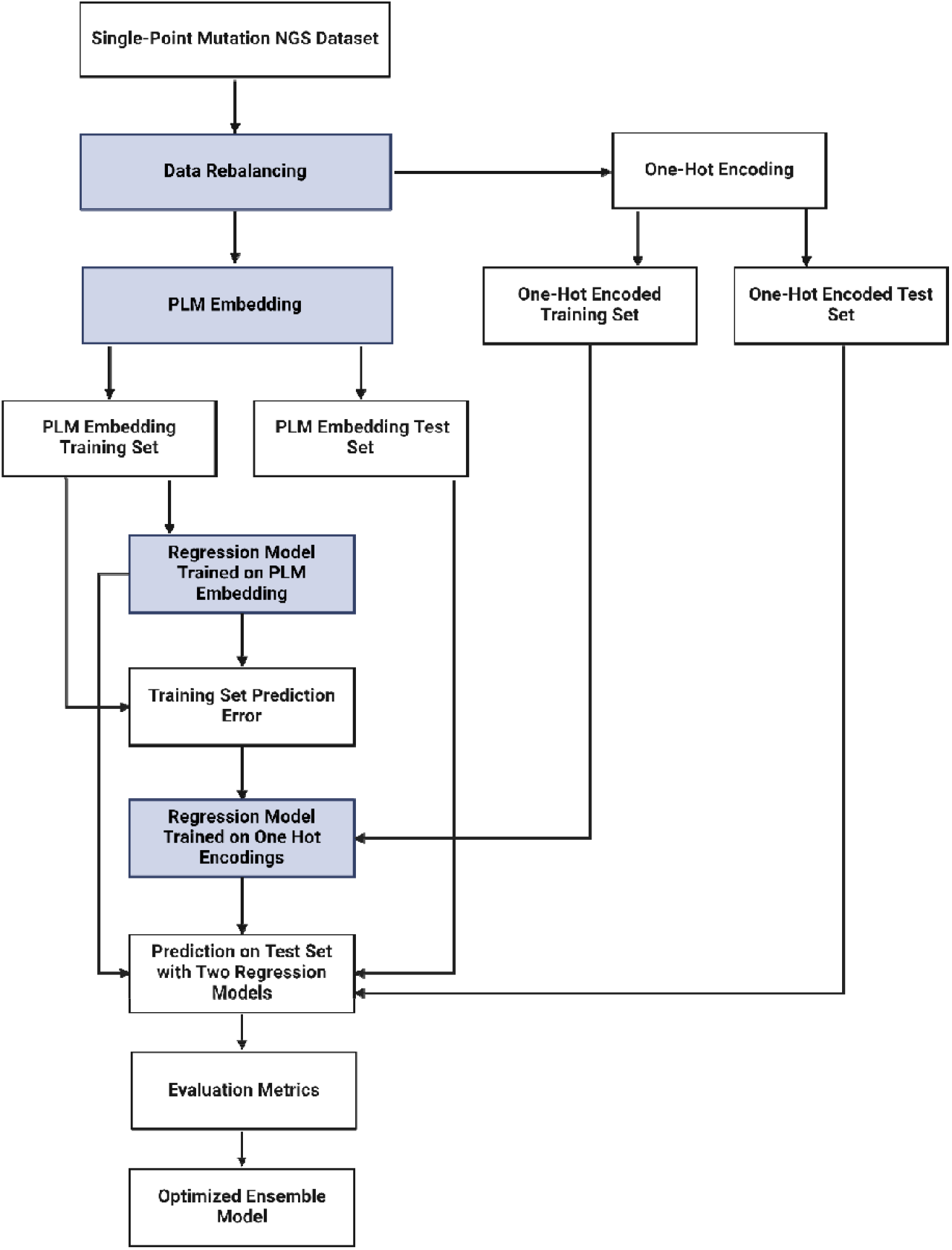
A Diagram of Steps Taken to Train CAP-PLM. Here, we highlight the general workflow of how CAP-PLM was trained. Highlighted in periwinkle are steps that one changes to improve performance of the CAP-PLM model on the evaluation metrics.

### Validation of model on additional datasets

To test the robustness and determine if the predictive power of CAP-PLM extends beyond the single point mutants data on which it was trained, we tested the method against two additional datasets, a library of 93 AAV2 mutants generated by site-directed mutagenesis of 48 approximately evenly spaced sites in the *cap* gene,(18) and a library of capsids generated by saturation mutagenesis of positions 561-588 of the AAV2 VP1 sequence.(3) These two datasets contain multiple-point mutations; therefore, we chose to validate with the multiple-point mutations that are not contained within the single amino acid mutations in our original training/test dataset.

For the first dataset, we analyzed two types of mutants: class A (n=13) mutants were fully competent and had titers comparable to wild-type AAV2, while class B mutants (n=8) were non-assembling. We found that the predicted fitness scores of the class B (non-assembling) mutants was significantly (KS-test, p<0.005) lower than those of the class A (fully competent) mutants (Figure 2A). Therefore, despite being trained on a single mutation dataset, CAP-PLM is robust enough to extrapolate to capsids harboring more than one mutation.

We then sought to test CAP-PLM in a much larger mutant dataset. Unlike the first small dataset,(18) containing only 23 mutants, we next explored a dataset(3) contains 201,426 mutants that were between 2 and 39 mutations away from wild-type AAV2 capsid sequence. We divided these mutants by number of mutations into viable or non-viable mutants as assigned by the original publication, predicted fitness scores for these mutants and compared viable and non-viable mutants (Figure 2B). We found the predicted fitness scores of non-viable mutants with 2-25 mutations were significantly lower than for viable mutants by KS-test (fdr corrected KS-test, p<.05), but the test statistic of the comparison decreases as the number of mutations increases (Figure 2C), suggesting that the prediction power of CAP-PLM decreases as the number of mutations increases. In conclusion, CAP-PLM is a flexible and extensible framework which is a strong predictor even on multi-mutant AAV2 fitness. Furthermore, CAP-PLM is generalizable to make predictions across the entire AAV VP1 sequence.

## Discussion

We have established a state-of-the-art ML model which accurately predicts the fitness of AAV2 capsid mutants from the amino acid sequence of VP1. Using this model, we can rapidly analyze far more sequences than can be screened using biological assays, filtering out inviable sequences, reducing costs, and improving efficiency in capsid library designs. As these dead mutants represent wasted library space, this increases the chance that a given library will contain variants with biological properties of interest.

We trained our model on a library of AAV2 mutants, the most comprehensive AAV capsid mutagenesis library available at the time. However, the approach is flexible and could provide a framework that could be extended to other clinically relevant AAV serotypes. This would require an initial investment in a single mutant library. This library could be leveraged to explore additional capsid properties, making such an investment potentially valuable. In addition, the model itself might be adapted or extended to predict additional biological properties, although this would likely require specific modifications in model tuning.

Our choice of model was driven by the application of natural language processing to biological problems. PLM has been shown to perform well on stability and function prediction, while ESM-2 performs well on structure prediction. However, other PLMs, such as Deep SequenceVAE(19) or EVmutation(20), have been shown to outperform PLM and ESM-1b(8), an earlier version of ESM-2, on fitness prediction for a variety of different proteins.(16) It is likely that different types of models encode different information about the proteins for usage in a downstream task. Therefore, it is possible that a different PLM might perform better if the model was adapted for the prediction of many different types of proteins.

In developing this model, we prioritized a lightweight model with reduced potential for overfitting. Therefore, we used a classical ML model rather than a neural network. This architecture is suitable for our relatively small dataset. However, it may not be scalable to larger training datasets, for prediction of more complicated capsid properties, or for prediction of multiple properties. A neural network-based architecture could potentially improve the scalability of the model and allow it to predict more complex capsid properties. Indeed, neural network-based architectures have been used to predict capsid assembly and to generate diverse viable sequence variant,(3,12) but these methods did not incorporate PLMs. The combination of a PLM that is trained off implicit fitness constraints present in known protein sequences, with site-specific fitness can outperform either method alone.

While this predictive model is useful on its own, combining it with a generative model would be a consequential leap forward in AAV engineering. At present, most AAV libraries designs focus on known regions of the AAV capsid sequence. However, a generative model could be used to design a capsid library agnostic to known AAV biology, expanding the sequence space that could be explored for potentially therapeutically useful capsids while ensuring that libraries are not dominated by non-assembling capsids. Furthermore, capsids generated by such a model might exceed the parental capsid in fitness which, in the long term, could contribute to lowering the COGS for AAV gene therapies, making them more affordable and accessible. Finally, capsids generated by such a model might elucidate the functions of previously unexplored regions of capsid genomes. The predictive model described herein is a critical first step towards solving key issues in AAV engineering.

## Methods

### Predictive Model Development (CAP-PLM)

We use the single point mutant data to train an ML model to predict the fitness of AAV2 mutants utilizing custom Python scripts. (21,22) Fitness is described in the same method as original publication.(14) Specific definitions can be found in Supplementa1 Equation 1.

After extracting the sequences and fitness scores from the training data, we embedded the sequences using PLM or ESM-2 and used these embeddings as the inputs for top model training. For PLM, we utilized a previously described implementation.(23) We did a randomized 80-20 training-test set split, setting 20% of the embeddings aside for validation of the top model. Using scikit-learn, we performed a grid search on K-Nearest Neighbors Regression, Support Vector Regression, and Random Forest Regression to see which top model produces the highest Spearman/Pearson correlation with the experimental data on the set aside test set. This same 80-20 split was used across all the later hyperparameter tuning procedures, with k-fold cross validation search across the training and test set showing little difference in the Pearson and Spearman correlation on the given test set.

## Validation of CAP-PLM

We validated CAP-PLM using a library of 23 mutants generated by site-directed mutagenesis for which individual infectious titer assays were available. We selected Class A mutants, which had production and infectious titers similar to WT AAV2 and Class B non-producing mutants and manually parsed the mutations from the original paper. We then used our model to predict their fitness scores and compared them using KS-test. Additional validation was performed using a library generated by saturation mutagenesis of positions 561-588 of the AAV2 VP1 sequence for which fitness was measured.(3) We compared predicted fitness to measured fitness using KS-test.

## Supporting information

Supplementary Figures and Equations

## Acknowledgements

We thank Edith Pfister, Ph.D. (Sanofi, Genomic Medicine Unit) for assistance with drafting and editing this manuscript. We gratefully acknowledge the use of the GMU Magellan platform, our Sanofi R&D compute platform, for facilitating data analysis and collaboration during the course of this research. Figures were created with BioRender.com.

