## Supplementary Figures and Equations for "Prediction of Adeno-Associated Virus Fitness with a Protein Language Based Machine Learning Model"

### Supplementary Information

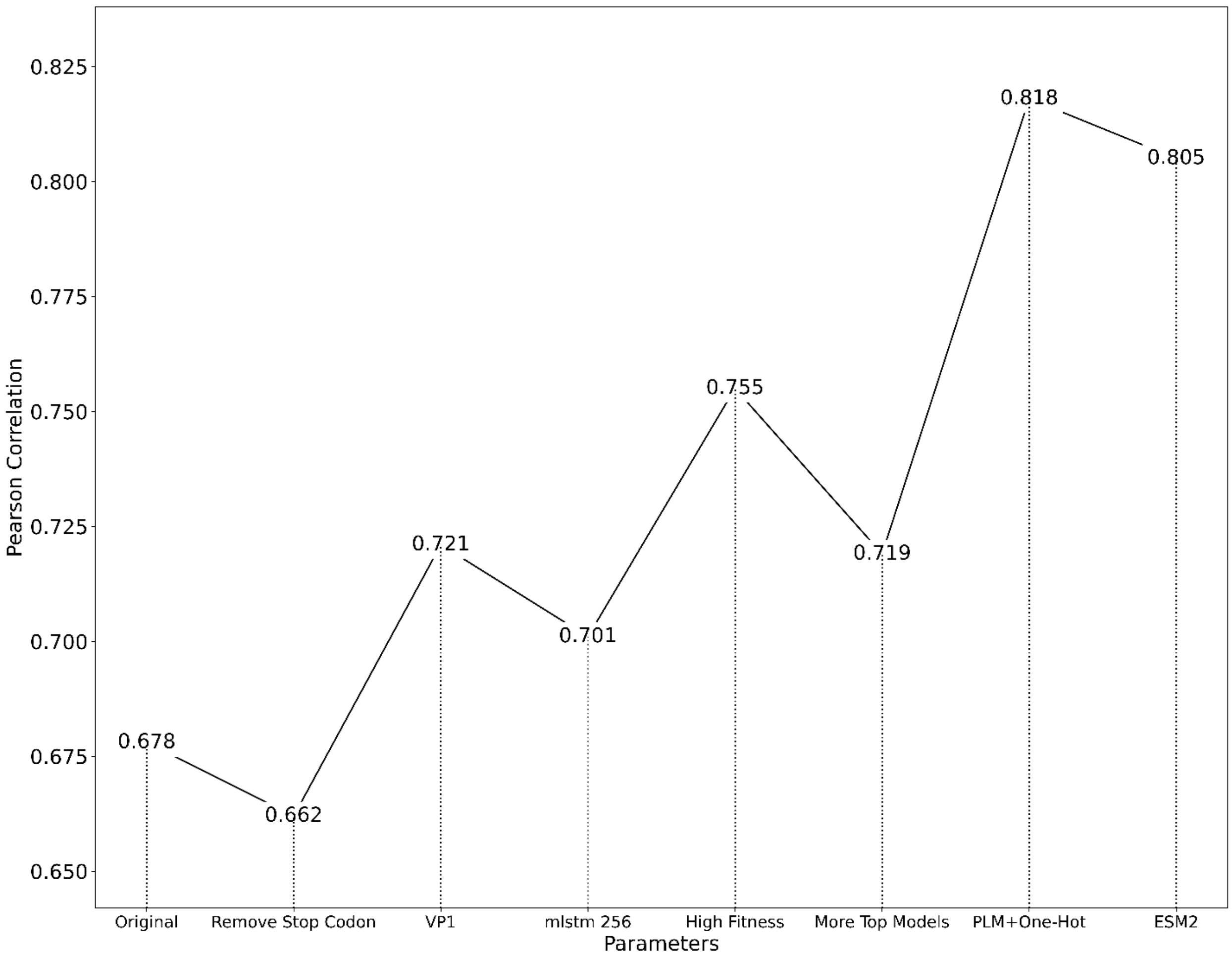


**Supplementary Figure 1. Parameters searched during model training.** Pearson correlation for each model against a held-out test set is plotted.

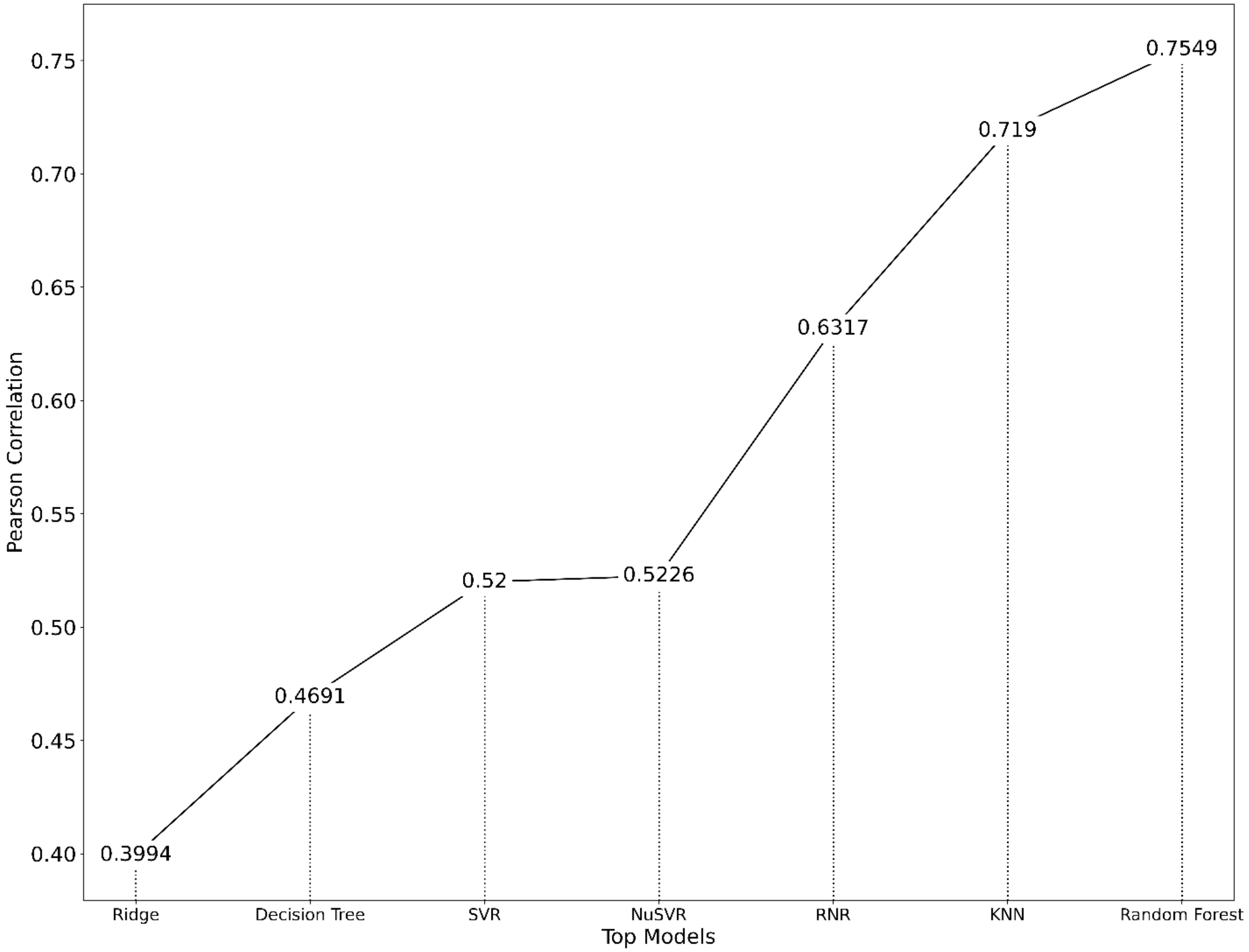


**Supplementary Figure 2. Comparison Between Different Top Regression Models.** Pearson correlation of model predictions upon the same held-out test set is plotted.

**Supplementary Table 1. Summary of Steps Taken in Predictive Model Training.**

| Training Steps | Description |
| --- | --- |
| Reformat | Reformat AAV capsid sequences and their fitness scores. |
| Data Rebalancing | Rebalance the training data to improve the performance of the regression model. |
| Embed | Embed the sequences using either ESM-2 or PLM for top-model training. |
| Train | Train using different top models to build a regression model. |
| Predict | Predict which AAV capsid sequences would have the highest selection weights and compare against the experimental data. |

**Supplementary Table 2. Specifics on Hyperparameters and Methods Investigated in Model Optimization.**

| Hyperparameter Categories | Specific Hyperparameters Tested |
|---|---|
| Embedding Models | Evolutionary Scale Modeling (ESM-2) |
| | PLM |
| Data Rebalancing | Remove stop codon sequences |
| | Rebalance data by removing sequences with low fitness scores |
| AAV Monomers | VP3 |
| | VP1 |
| Embedding Dimensions | 1900 |
| | 256 |
| Regression Models | Ridge |
| | K-Nearest Neighbors/Radius Nearest neighbors |
| | Support Vector Regression/Nu Support Vector Regression |
| | Random Forest Regression/Decision Tree Regression |
| PLM Embeddings +One-Hot Encoding | Train a separate top model with one-encoded representations and the error of the predictions upon the original regression model. |

$$\frac{\frac{\boldsymbol{Plasmid1ReadCounts_n}}{\boldsymbol{TotalPlasmid1ReadCounts}} + \frac{\boldsymbol{Plasmid2ReadCounts_n}}{\boldsymbol{TotalPlasmid2ReadCounts}}}{\mathbf{2}} = \boldsymbol{NormalizedPlasmidReadCounts_n}$$

$$\frac{\frac{\boldsymbol{Viral1ReadCounts_n}}{\boldsymbol{TotalViral1ReadCounts}} + \frac{\boldsymbol{Viral2ReadCounts_n}}{\boldsymbol{TotalViral2ReadCounts}}}{\mathbf{6}} = \boldsymbol{NormalizedViralReadCounts_n}$$

$$\frac{\boldsymbol{NormalizedPlasmidReadCounts_n}}{\boldsymbol{NormalizedViralReadCounts_n}} = \boldsymbol{FvFp_n}(\boldsymbol{Manufacturability})$$

$$\frac{\boldsymbol{FvFp_n}}{\boldsymbol{FvFp_{AAV2}}} = \boldsymbol{NormalizedFvFp_n}$$

$$\log_2(\boldsymbol{NormalizedFvFp_n}) = FitnessScore$$

**Supplementary Equation 1. Equations used to calculate fitness score of AAV mutants.** *n* refers to number of distinct AAV capsids, with each AAV capsid having a unique barcode. Plasmid and viral read counts are normalized to account for variations in sequencing depth.
